# ACE2 polymorphisms and individual susceptibility to SARS-CoV-2 infection: insights from an *in silico* study

**DOI:** 10.1101/2020.04.23.057042

**Authors:** Matteo Calcagnile, Patricia Forgez, Antonio Iannelli, Cecilia Bucci, Marco Alifano, Pietro Alifano

## Abstract

The current SARS covid-19 epidemic spread appears to be influenced by ethnical, geographical and sex-related factors that may involve genetic susceptibility to diseases. Similar to SARS-CoV, SARS-CoV-2 exploits angiotensin-converting enzyme 2 (ACE2) as a receptor to invade cells, notably type II alveolar epithelial cells. Importantly, ACE2 gene is highly polymorphic. Here we have used *in silico* tools to analyze the possible impact of ACE2 single-nucleotide polymorphisms (SNPs) on the interaction with SARS-CoV-2 spike glycoprotein. We found that S19P (common in African people) and K26R (common in European people) were, among the most diffused SNPs worldwide, the only two SNPs that were able to potentially affect the interaction of ACE2 with SARS-CoV-2 spike. FireDock simulations demonstrated that while S19P may decrease, K26R might increase the ACE2 affinity for SARS-CoV-2 Spike. This finding suggests that the S19P may genetically protect, and K26R may predispose to more severe SARS-CoV-2 disease.

## Introduction

In principle, any new infectious agent that challenges a totally susceptible population with little or no immunity against it is able to totally infect the population causing pandemics. Pandemics rapidly spread affecting a large part of people causing plenty of deaths with significant social disruption and economic loss. However, if we look at the even worst pandemics in the human history we can realize that ethnic and geographical differences in the susceptibility to disease actually exist, in spite of the infectious sources and transmission routes that are the same for all individuals^1^. The current SARS covid-19 (a shortened form of “coronavirus disease of 2019”) epidemic spread appears to be similarly influenced by ethnical and geographical factors. After its initial spread in China, the pandemic is now progressing at an accelerating rate in Western Europe and the United States of America^2^. In these regions, the causative agent, the severe acute respiratory syndrome corona virus −2 (SARS-CoV-2) is spreading incredibly quickly between people, due to its newness – no one on earth has immunity to SARS Covid-19 – and transmission route. Yet, in the other regions of the world, the kinetics of diffusion and mortality seem less impressive, although the world has become highly interconnected as a result of a huge growth in trades and travels^2^.

A multitude of factors may concur to explain the ethnic and geographical differences in pandemic progression and severity, including cultural, social and economic inequality, as well as health care system organization, and climate also. Mostly, considerable individual differences in genetic susceptibility to diseases may be involved^3^. Genomic predisposition is a major concept in modern medicine, and understanding of molecular bases of genetic predisposition can help to find prevention and treatment strategies for the corresponding diseases. In the SARS Covid-19, even subtle inter-individual genetic differences may affect both the SARS-CoV-2 viral life cycle and the host innate and acquired immune response.

SARS-CoV-2 is an enveloped positive-stranded RNA virus that replicates in the cytoplasm, and uses envelope spike projections as a key to enter human airway cells^4^. In coronaviruses spike glycoproteins, which forms homotrimers protruding from the viral surface, are a primary determinant of cell tropism, pathogenesis, and host interspecies transmission. Spike glycoproteins comprise two major functional domains: an N-terminal domain (S1) for binding to the host cell receptor, and a C-terminal domain (S2) that is responsible for fusion of the viral and cellular membranes^5^.

Following the interaction with the host receptor, internalization of viral particles into the host cells is accomplished by complex mechanisms that culminate with the activation of fusogenic activity of spike, as a consequence of major conformational changes that, in general, may be triggered by receptor binding, low pH exposure and proteolytic activation^5^.

In some coronaviruses spike glycoproteins are cleaved by furin, a Golgi-resident protease, at the boundary between S1 and S2 domains, and the resulting S1 and S2 subunits remain non-covalently bound in the prefusion conformation with important consequences on fusogenicity^5^. Notably, at variance with SARS-CoV and other SARS-like CoV Spike glycoproteins, SARS-CoV-2 Spike glycoprotein contain a furin cleavage site at the S1/S2 boundary, which is cleaved during viral biogenesis ^6^, and may affect the major entry route of viruses into the host cell^5^.

Productive entry of coronaviruses that harbor non-cleaved Spike glycoproteins (such as SARS-CoV) rely on endosomal proteases suggesting that this entry is accomplished by hijacking the host endocytic machinery^5^. Indeed, it has been reported that SARS-CoV infection is inhibited by lysomotropic agents because of the inhibition of the low-pH-activated protease cathepsin L^7^. However, SARS-CoV is also able to fuse directly to the cell membrane in the presence of relevant exogenous proteases, and this entry route is believed to be much more efficient compared to the endocytic route^8^. In fact, proteases from the respiratory tract such as those belonging to the transmembrane protease/serine subfamily (TMPRSS), TMPRSS2 or HAT (TMPRSS11d) are able to induce SARS-CoV spike glycoprotein fusogenic activity^9,10,11,12^. The first cleavage at the S1-S2 boundary (R667) facilitates the second cleavage at position R797 releasing the fusogenic S2’ sub-domain^5^. On the other hand, there is also evidence that cleavage of the ACE2 C-terminal segment by TMPRSS2 can enhance spike glycoprotein-driven viral entry^13^. Notably, it has been very recently demonstrated that also SARS-CoV-2 cell entry depends on TMPRSS2, and is blocked by protease inhibitors^14^.

SARS-CoV-2 and respiratory syndrome corona virus (SARS-CoV) Spike proteins share very high phylogenetic similarities (99%), and, indeed, both viruses exploit the same human cell receptor namely angiotensin-converting enzyme 2 (ACE2), a transmembrane enzyme whose expression dominates on lung alveolar epithelial cells^6,15,16^. This receptor is an 805-amino acid long captopril-insensitive carboxypeptidase with a 17-amino acids N-terminal signal peptide and a C-terminal membrane anchor. It catalyzes the cleavage of angiotensin I into angiotensin 1-9, and of angiotensin II into the vasodilator angiotensin 1-7, thus playing a key role in systemic blood pressure homeostasis, counterbalancing the vasoconstrictive action of angiotensin II, which is generated by cleavage of angiotensin I catalyzed by ACE^17^ Although ACE2 mRNA is expressed ubiquitously, ACE2 protein expression dominates on lung alveolar epithelial cells, enterocytes, arterial and venous endothelial cells, and arterial smooth muscle cells^18^.

There is evidence that ACE2 may serve as a chaperone for membrane trafficking of an amino acid transporter B0AT1 (also known as SLC6A19), which mediates the uptake of neutral amino acids into intestinal cells in a sodium dependent manner^19^. Recently, 2.9 Å resolution cryo-EM structure of full-length human ACE2 in complex with B0AT1 was presented, and structural modelling suggests that the ACE2-B0AT1 can bind two spike glycoproteins simultaneously^20,21^. It has been hypothesized that the presence of B0AT1 may block the access of TMPRSS2 to the cutting site on ACE2^20,21^. B0AT1 (also known as SLC6A19) is expressed with high variability in normal human lung tissues, as shown by analysis of data available in Oncomine from the work by Weiss et al^22^.

Notably, a wide range of genetic polymorphic variation characterizes the ACE2 gene, which maps on the X chromosome, and some polymorphisms have been significantly associated with the occurrence of arterial hypertension, diabetes mellitus, cerebral stroke, coronary artery disease, heart septal wall thickness and ventricular hypertrophy^23,24,25^. The association between ACE2 polymorphisms and blood pressure responses to the cold pressor test led to the hypothesis that the different polymorphism distribution worldwide may be the consequence of genetic adaptation to different climatic conditions^25,26^. In this study we have used a combination of *in silico* tools to analyze the possible impact of ACE2 single-nucleotide polymorphisms (SNPs) on the interaction with SARS-CoV-2 Spike glycoprotein. Results seem to suggest that ACE2 polymorphism can contribute to ethnic and geographical differences in SARS COVID-19 spreading across the world.

## Results

### 3D Protein Data Bank (PDB) models and *in silico* screening of ACE2 SNPs affecting binding with coronavirus Spike proteins

All simulations were carried out using different 3D Protein Data Bank (PDB) X-ray cristallography models from RCBS (https://www.rcsb.org/): 2AJF (SARS-CoV Spike Receptor Binding Domain [RBD] /ACE2 complex)^27^, 6LZG (SARS-CoV-2 Spike RBD /ACE2 complex)^28^, 6M0J (SARS-CoV-2 Spike RBD /ACE2 complex)^29^, 6VW1 (chimeric SARS-CoV/SARS-CoV-2 Spike RBD /ACE2 complex)^30^, and 6M17 (SARS-CoV-2 Spike RBD /ACE2/B0AT1 complex)^20,21^ (Fig. 1a). ClustalO alignments of human ACE2 amino acid sequences, and SARS-CoV, SARS-CoV-2 and chimeric SARS-CoV/SARS-CoV-2 spike RBDs used for X-ray cristallography models are reported in Supplementary Fig. 1. 6VW1 was developed with a chimeric RBD to facilitate crystallization, by using the core from SARS-CoV RBD as the crystallization scaffold and the Receptor Binding Motif (RBM) from SARS-CoV-2 as the functionally relevant unit. Nevertheless, the structures of chimeric SARS-CoV/SARS-CoV-2 Spike RBD /ACE2 complex of 6VW1 and SARS-CoV-2 Spike RBD /ACE2 complex of 6M0J, particularly in RBM region, were highly similar^30^.

**Fig. 1.**
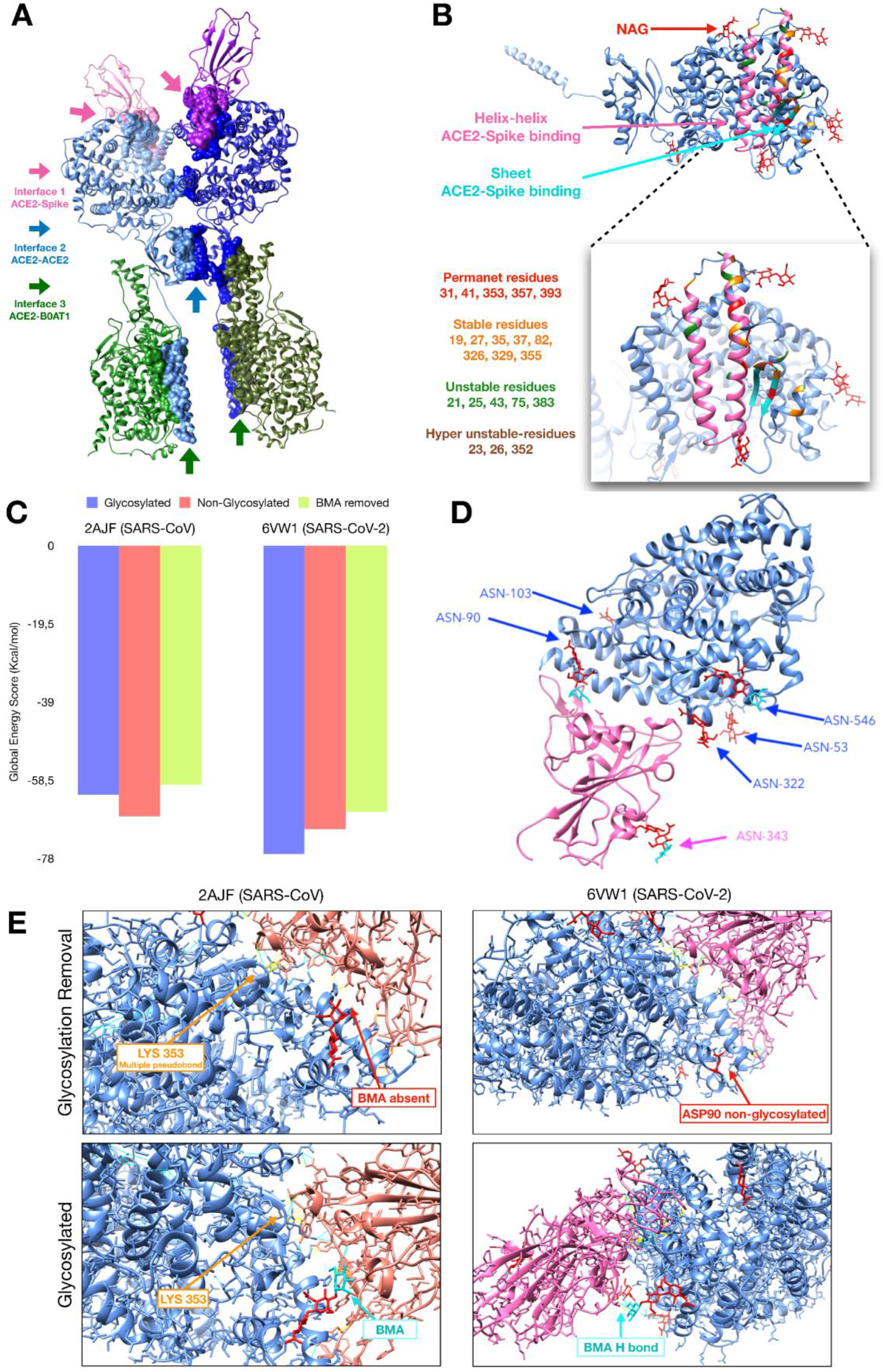
3D models of SARS-CoV-2 Spike/ACE2 complex, and effects of ACE2 glysosylation on binding interfaces. **a** 3D model of SARS-CoV-2 Spike/ACE2/B0AT1 complex. The arrows indicate the principal binding interfaces between all proteins involved in this complex. **b** Domains and ACE2 amino acid residues involved the interaction with Spike. “Permanent” (red), “stable” (orange), “unstable” (green), and “hyper-unstable” residues are indicated. **c** FireDock was used to estimate ΔG values by using 2AJF (SARS-CoV Spike/ACE2 complex) and 6VW1 (SARS-CoV-2 Spike/ACE2 complex) after removal of the glycosydic chains (red bars) or the terminal beta-mannose (BMA) (green bars). **d** 3D 6VW1 model with highlighting glycosydic chains (red, N-acetyl-glucosamine [NAG]; cyan, BMA). **e** Predicted effects of removal of the glycosydic chains or the BMA in 2AJF and 6VW1 models. In 6VW1, the BMA forms two H-bonds and one pseudobond, while these bonds are lost in non-glycosylated model. In contrast, in the 2AJF, the glycosydic chain forms only one H-bond and, after removing BMA the Thr-41 has more grads of binding.

In all models, similar to SARS-CoV RBM, SARS-CoV-2 RBM forms a concave surface that houses a convexity formed by two helices on the exposed surface of ACE2. Strong network of H-bond and salt bridge interactions mediate the receptor-ligand binding. Global energy and several distinctive features of the 3D models with and without glycosylation are reported in Supplementary Table 1. Contact residues are classified as: “permanent” (predicted as binding residues in all 10 models), “stable” (predicted as binding residues in 6 or 7 out 10 models), “unstable” (predicted as binding residues in 6 or 7 out 10 models), “hyper-unstable” (1 or 2 models out of 10).

Using the 3D PDB models and EVOEF^31,32^ on SSIPe web-server (https://zhanglab.ccmb.med.umich.edu/SSIPe/) we screened the entire list of 301 ACE2 SNPs causing missense mutations from the dbSNP and UNIPROT database to identify possible amino acid substitutions that may affect binding interfaces (Supplementary Table 2). SSIPe^33^ was used to estimate ΔΔG values associated with each amino acidic substitution, and to generate models of ACE2 polymorphic variants. The list of the amino acid substitutions that may affect the ACE2/Spike, ACE2/B0AT1 and ACE2/ACE2 interfaces is reported in Supplementary Table 3. Twenty-seven substitutions were predicted to influence the ACE2/Spike interface in at least one of the different 3D PDB models. Fifteen and seventeen were predicted to affect the ACE2/B0AT1 and ACE2/ACE2 interfaces, respectively. Some residues, which are described in UNIPROT database (https://www.uniprot.org/uniprot/Q9BYF1) as important for the interaction between spike and ACE2, are permanent contact residues (predicted as binding) but all of these are non-polymorphic (Fig. 1b). In contrast, polymorphic residues are stable, unstable or hyper-unstable. A list of 18 SNPs from dbSNP (S19P, I21T, I21V, E23K, A25T, K26E, K26R, T27A, E35D, E35K, E37K, S43R, E75G, M82I, G326E, E329G, G352V; D355N), which were predicted to affect the ACE2/Spike interface, was used for further analysis.

### SNPs possibly affecting ACE2 glysosylation

Supplementary Table 1 illustrates amino acid glycosylation sites, and structure of the glycosidic chains as inferred from different ACE2/Spike complex PDB models. Putative polymorphic sites (Q60R, N103H, N546D, N546S) from dbSNP database that may affect ACE2 glycosylation are also reported. One of these amino acid variations, N546D, is rather common in South Asia (Supplementary Table 4).

FireDock^34^ was used to estimate the effects of removal of glycosidic residues or chains on ACE2 interaction with SARS-CoV-2 Spike RBD by calculating ΔG values. The data indicated that removal of glycosidic chains results in either an increased or a decreased ΔG values, depending on the PDB model (Fig. 1c). In particular, removal of glycosidic moieties apparently strengthened the ACE2/Spike interaction in SARS-CoV Spike/ACE2 in the 2AJF model, while it appeared to weaken the interaction between SARS-CoV-2 Spike and ACE2 in the 6VW1 model. In both cases, the effect was mostly due to removal of the terminal beta-mannose (BMA) (Fig. 1c), which was predicted to decorate a glycosidic chain attached to aspartic amino acid residue at position 90 that maps in a helix that is involved in the interaction with Spike, as shown in Fig. 1d. Noteworthy, in the 6VW1 model, BMA is involved in two H-bonds and one pseudo-bond (Fig. 1e), and these bonds are lost in non-glycosylated models. In contrast, in the 2AJF model, the BMA forms only one H-bond, and after removal of terminal BMA, the Thr-41 acquires more grads for binding thereby strengthening the interaction with ACE2. These results seem to suggest that ACE2 glycosylation may play a different role in modulating the interaction with SARS-CoV Spike and SARS-CoV-2 Spike.

### Selected ACE2 SNPs affecting binding interfaces

FireDock^34^ was used to estimate the effects of selected ACE2 SNPs on interaction with SARS-CoV Spike RBD (2AJF model) and SARS-CoV-2 Spike RBD (6M0J and 6WV1 models). Selection was based on both frequency of these SNPs worldwide, and predicted effects on ACE2 binding interfaces. Screening was preceded by correlation analysis of SNPs data from different databases (Supplementary Fig. 2). Network plot (Fig. 2a) and Non-Metric Multidimensional Scaling (NM-MDS, Bray-Curtis index) (Fig. 2b) of the most diffused SNPs demonstrated that S19P and K26R were, among the most diffused SNPs worldwide, the only two SNPs that were able to potentially affect the interaction of ACE2 with SARS-CoV Spike and SARS-CoV-2 Spike (Supplementary Table 3). In particular, the S19P SNP is rather common in African people with a frequency about 0.3%, while K26R SNP is frequent in European people with a frequency about 0.5% (Supplementary Table 4).

**Fig. 2.**
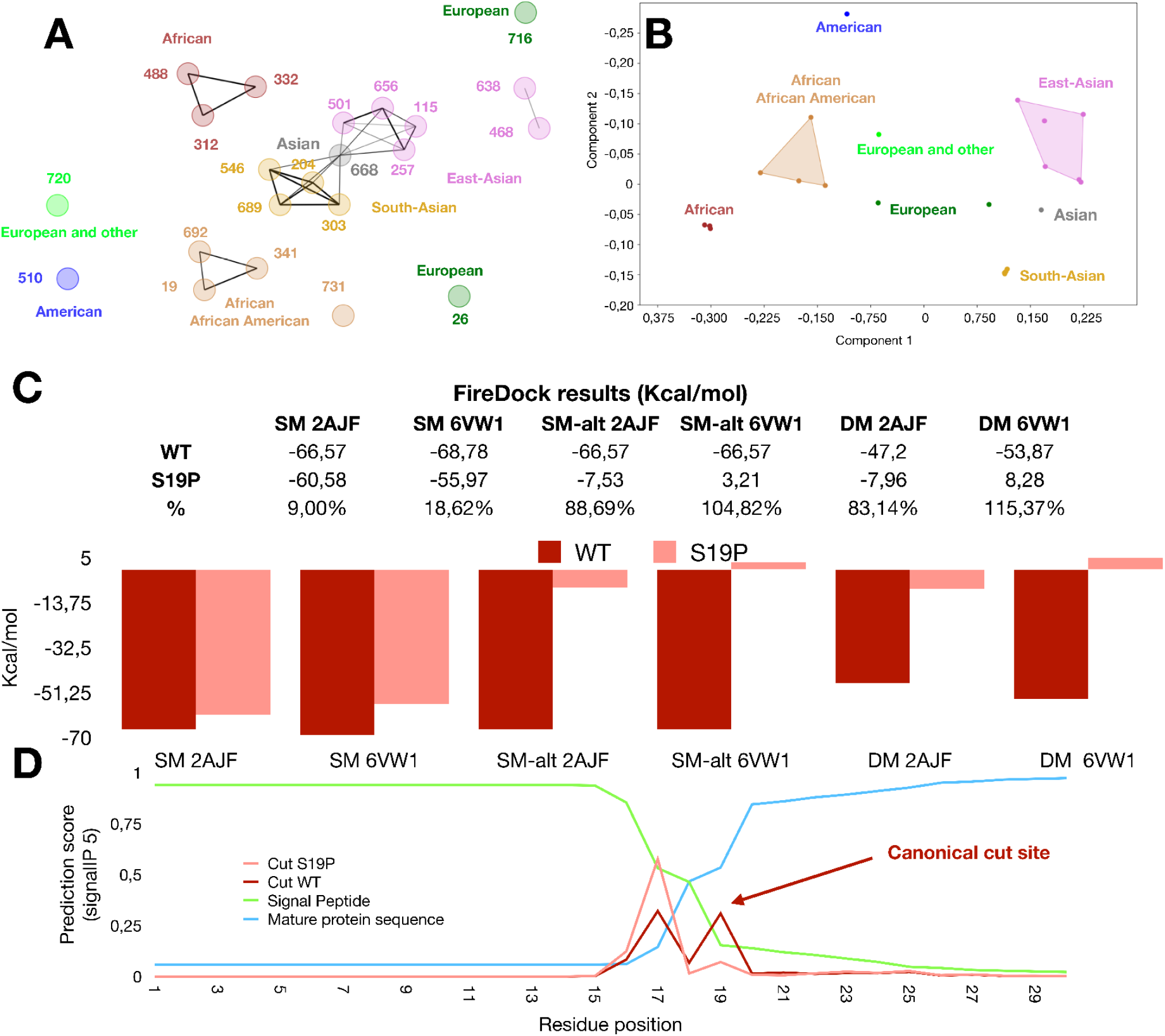
Selected ACE2 SNPs affecting binding interfaces: S19P. **a** Network plot of the most diffused ACE2 SNPs causing missenses worldwide. **b** Non-Metric Multidimensional Scaling (NM-MDS, Bray-Curtis index) the most diffused ACE2 SNPs. **c** FireDock results predicting the effects of ACE2 S19P amino acid replacement on ACE2/Spike interaction in 2AJF (SARS-CoV Spike/ACE2), and 6VW1 (chimeric SARS-CoV/SARS-CoV-2 Spike RBD /ACE2) models. Results predicting the effects of ACE2 S19P with the alternative (ALT) N-terminal cleavage site **(d)** are also shown in either 2AJF and 6VW1 static models (SM-alt), or2AJF and 6VW1 dynamic (docking) models (DM). **d** Predicted effect of ACE2 S19P on ACE2 N-terminal cleavage site.

FireDock^34^ results indicated that the S19P substitution decreased the affinity of ACE2 with Spike in 2AJF and 6VW1 models (Fig. 2c) and similar results were obtained with all other models. Moreover, this amino acid substitution seems also to affect the ACE2 N-terminal cleavage site (Fig. 2d), and when FireDock^34^ simulations were carried out on ACE2 with the alternative cleavage site, the effects of S19P SNP was much more impressive (Fig. 2 c).

In contrast, the K26R and the less common K26E substitutions appeared to increase the affinity of ACE2 with SARS-CoV-2 Spike (2AJF model), and slightly decrease the affinity of ACE2 with SARS-CoV Spike (6VW1 and 6M17) models (Fig. 3a). As 6VW1 was generated with a chimeric SARS-CoV/SARS-CoV-2 Spike, to support our results we performed an additional simulation by challenging the ACE2 structure from 6VW1 with the Spike structures that were generated by the different models (Fig. 3b), and the results confirmed those shown in Fig. 3a. Noteworthy, the receptor-ligand interactions was much weaker in 6M17 (SARS-CoV-2 Spike RBD /ACE2/B0AT1 complex) with respect to the other models, confirming an inhibitory function of B0AT1. However, in this model, at lower energy values, the effects of K26R/E substitutions were much more evident. FireDock^34,35^ simulations indicate that such an increased affinity between K26R ACE2 and SARS-CoV-2 Spike could be due to an increased number of H-bond and/or pseudo-bonds around Glu-35, Met-82 and Lys-353 (Fig. 3c). Based on this result, the K26R and K26E could genetically predispose to more severe SARS-CoV-2 disease. In addition, several ACE2 SNPs were predicted to affect ACE2/ACE2 homo-dimerization (E668K, N638S, R716H, R710H) or ACE2/B0AT1 interaction (L731F) (Supplementary Fig. 3).

**Fig. 3.**
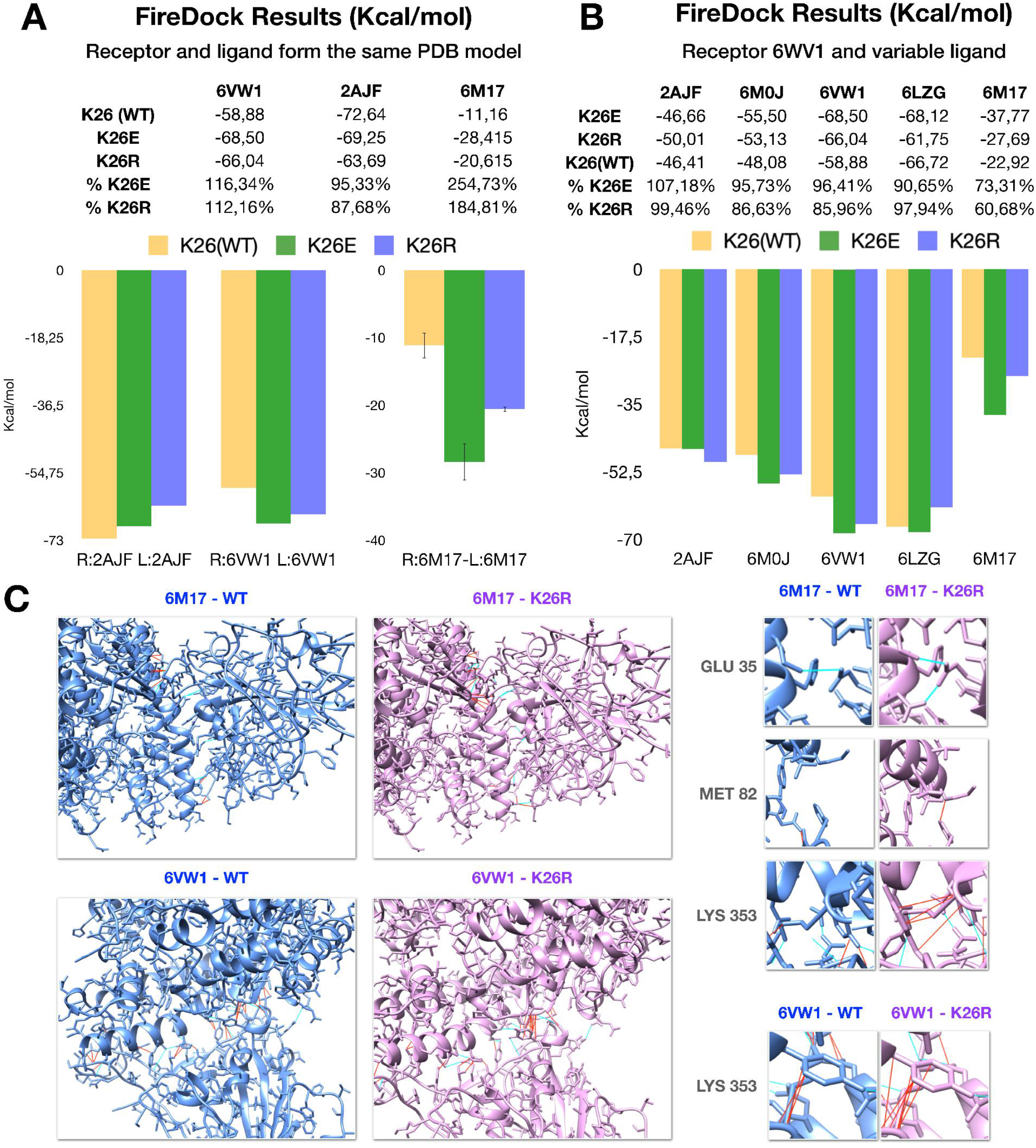
Selected ACE2 SNPs affecting binding interfaces: K26R/E. **a** FireDock results predicting the effects of ACE2 K26R and K26E amino acid replacements on ACE2/Spike interaction in 2AJF (SARS-CoV Spike/ACE2), and 6VW1 (chimeric SARS-CoV/SARS-CoV-2 Spike RBD /ACE2) and 6M17 (SARS-CoV-2 Spike RBD /ACE2/B0AT1). **b** FireDock results that were obtained by challenging the ACE2 structure from 6VW1 with the Spike structures that were generated by the different models as shown in the bottom of the histogram. **c** Possible effects of K26R substitutions on H-bonds (cyan) and pseudo-bonds (red) from 6M17 (upper images) and 6VW1 (upper panels) models as illustrated (larger panels: left, wild type ACE2; right, K26R ACE2). Smaller panels on the right are magnifications of regions the respective panels on the left as indicated.

### Dynamic models of ACE2: fluctuation, deformation and chaperone requirement for correct topology maintenance

Dynamut^36^ was used to analyze dynamic features of ACE2 receptor including fluctuation and deformation. Dynamut^36^ calculated fluctuation and deformation scores for each ACE2 amino acid residue position (Fig. 4a) thus providing dynamic features of the analyzed protein. The model shows that ACE2 (computed by using 3D PDB model 6M17 as input file) is characterized by a high deformation tract that is located immediately upstream of the transmembrane domain (Fig. 4b), whereas the C-terminal tail is characterized by high fluctuation (Fig. 4c).

**Fig. 4.**
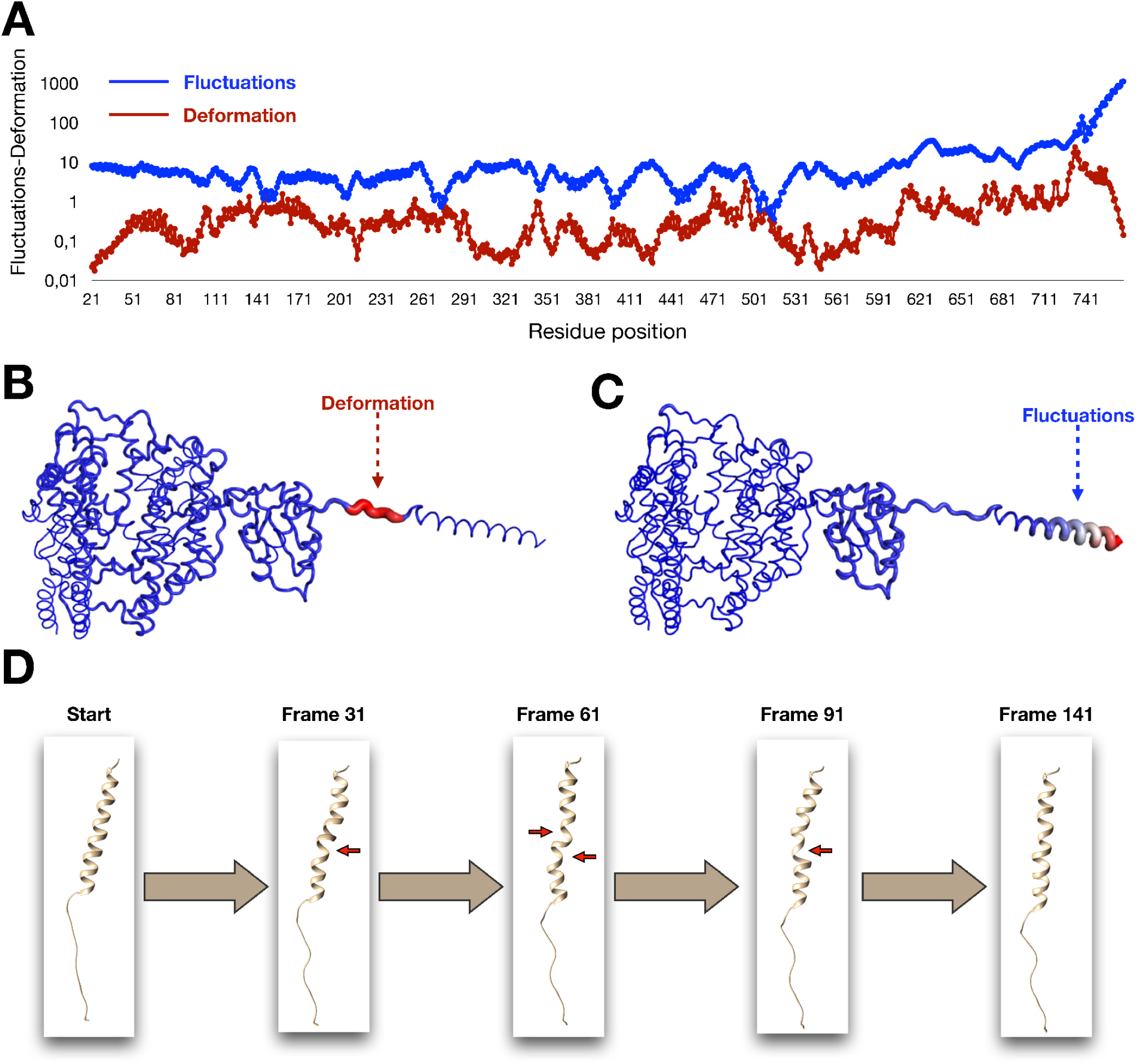
Dynamut models of ACE2. **a** Score for fluctuation and deformation of ACE2 calculated by Dynamut. **b** ACE2 region that is mostly subject to deformation. **c** ACE2 region that is mostly subject to fluctuation. D) Distortion of the hydrophobic transmembrane domain during Molecular Dynamic Simulation.

CHARMM-GUI^37,38^ and VMD/NAMD^39,40^ tools were used to model the interaction of the ACE2 transmembrane domain with the phospholipid membrane. CHARMM-GUI was used to build the model of the phospholipid membrane embedding a single chain of ACE2, while VMD/NAMD tools were used to perform Molecular Dynamics Simulation. Frames shown in Fig. 4d seem to suggest that the hydrophobic domain alone is highly unstable in the membrane confirming that a chaperone is required for correct topology maintenance. This function was assigned to the moonlighting amino acid transporter B0AT1^19^.

To investigate dynamic properties of ACE2 globular head, the trans-membrane helix and conserved domains were firstly mapped on a 3D structure. Then, Dynamut^36^ simulation was carried out on ACE2 by using 6WV1 PDB model (without the transmembrane domain) (Fig. 5ab). Results indicate that some residues of the ACE2 interface, which are involved in the interaction with SARS-CoV-2 Spike glycoprotein can actually fluctuate (Fig. 5cd).

**Fig. 5.**
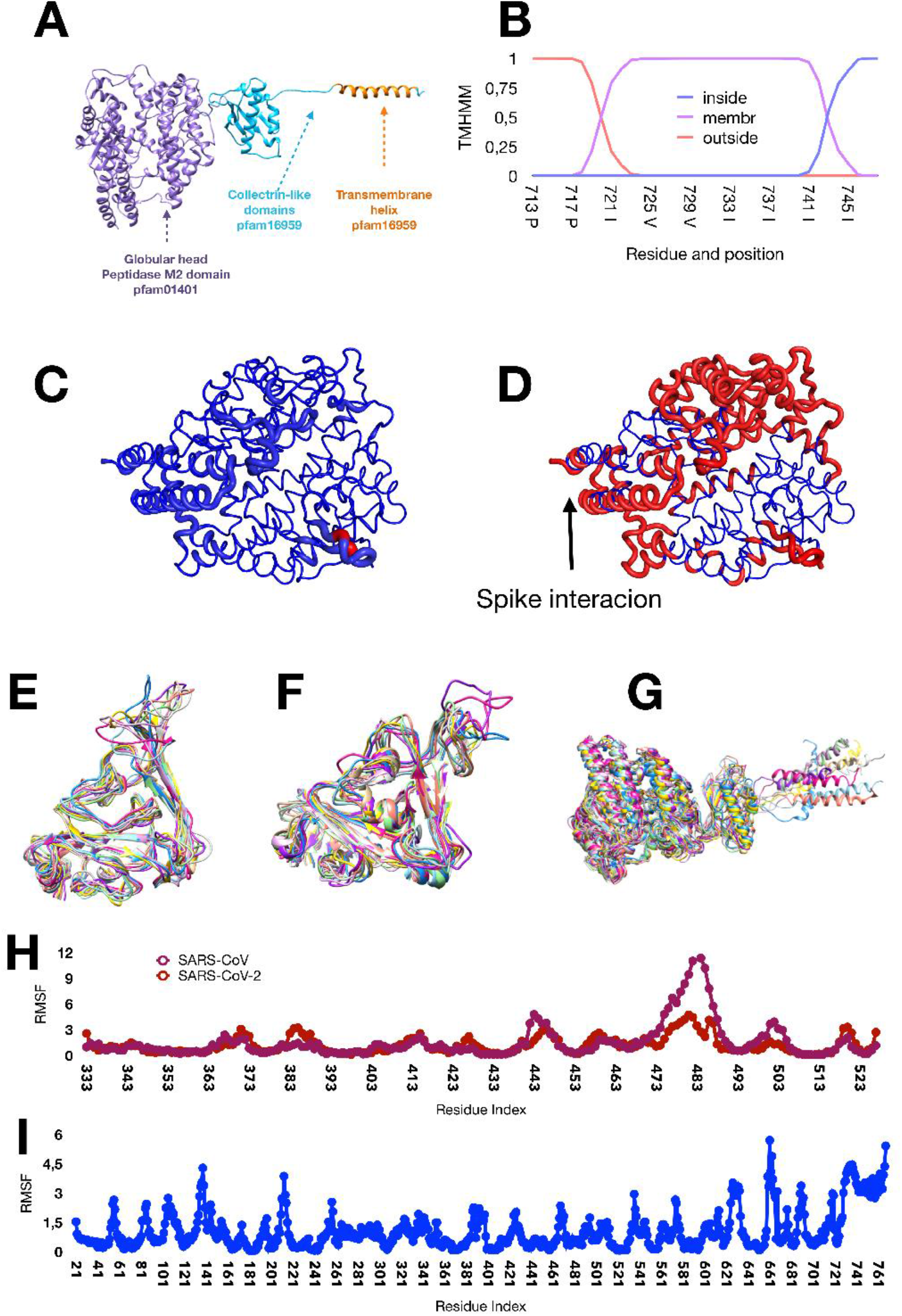
Structural and dynamic features of ACE2, SARS-CoV-2 and SARS-CoV Spike proteins as inferred by Dynamut and CABSflex. **a** Conserved domains of ACE2. **b** Transmembrane region of ACE2 identified by TMHMM. **c-d** Dynamut simulations of ACE2 structure by using the 6WV1 PDB model (without the transmembrane domain) showing that some residues of the ACE2 interface that bind the Spike protein can fluctuate. **e-g** CABSflex simulations of SARS-CoV (**e**), SARS-CoV-2 Spike (**f**), and ACE2 (**g**). **h** CABSflex fluctuation scores of SARS-CoV Spike and SARS-CoV-2 Spike. **i** CABSflex fluctuation scores of ACE2.

Dynamic properties of SARS-CoV-2 and SARS-CoV spike proteins were also investigated. CABS-flex^41^ was used to this purpose, also to compare fluctuation scores of SARS-CoV-2 and SARS-CoV Spike RBD (receptor binding domain) of spike and ACE2. Fig. 5ef depicts the CABS flex model of SARS-CoV and SARS-CoV-2 Spike, respectively while Fig. 5g illustrates the model of ACE2. CABS-flex model (Fig. 5ef) and fluctuation scores indicate that SARS-CoV Spike is characterized by higher flexibility of in a coiled-coil domain, with respect to SARS-CoV-2 Spike (Fig. 5h). This difference may affect the interaction with ACE2.

Variation of the distance between the amino acid residues involved in ACE2 binding interfaces were then analyzed by Molecular Dynamics Simulation, a computer simulation method for analyzing the physical movements of atoms and molecules. Supplementary Fig. 4a and 4c show, respectively, oscillation plot (in ångström) and variance of distance between the amino acid residues in the two helices that are involved in the SARS-CoV-2 Spike interaction, while Supplementary Fig. 4b illustrates 3D images of regions containing the amino acid residues of Supplementary Fig. 4a. Oscillation plots and variance of the distance of amino acid residues between the two helices and beta-sheet, and between the residues of the beta-sheet are illustrated in Supplementary Fig. 4d and 4f, and Supplementary Fig. 4e and 4g, respectively. Overall, the data indicate that, although the two helices in the binding interface with Spike protein form a compact structure, some residue can actually oscillate. Permanent amino acid residues (41-31) exhibit small oscillation by Molecular Dynamics Simulation. In contrast, stable or unstable residues exhibit a different behavior. The residue 21 occupies the first position in the model used (PDB 6M17) for simulation, and, as a consequence, it exhibits the highest oscillation degree. Interestingly, the distance between the amino acid residues 26-36 and 72-82 shows high variation indicating that Met-82 and Lys-26 may be characterized by considerable degree of freedom.

### ACE2 SNPs analyzed by dynamic models: structural effects on binding interfaces

Dynamut36 and ENCoM^42^ were used to compare dynamic features of ACE2 and ACE2 polymorphic variants. Both these tools predict ΔΔG values associated with single amino acid substitution; ENCoM^42^ also predicts ΔΔE values. The Dynamut^36^ and ENCoM^42^ outputs were used to generate ordination plots (PCA) by PAST to evaluate overall results (Fig. 6a, Supplementary Fig. 5 and Supplementary Table 7).

**Fig. 6.**
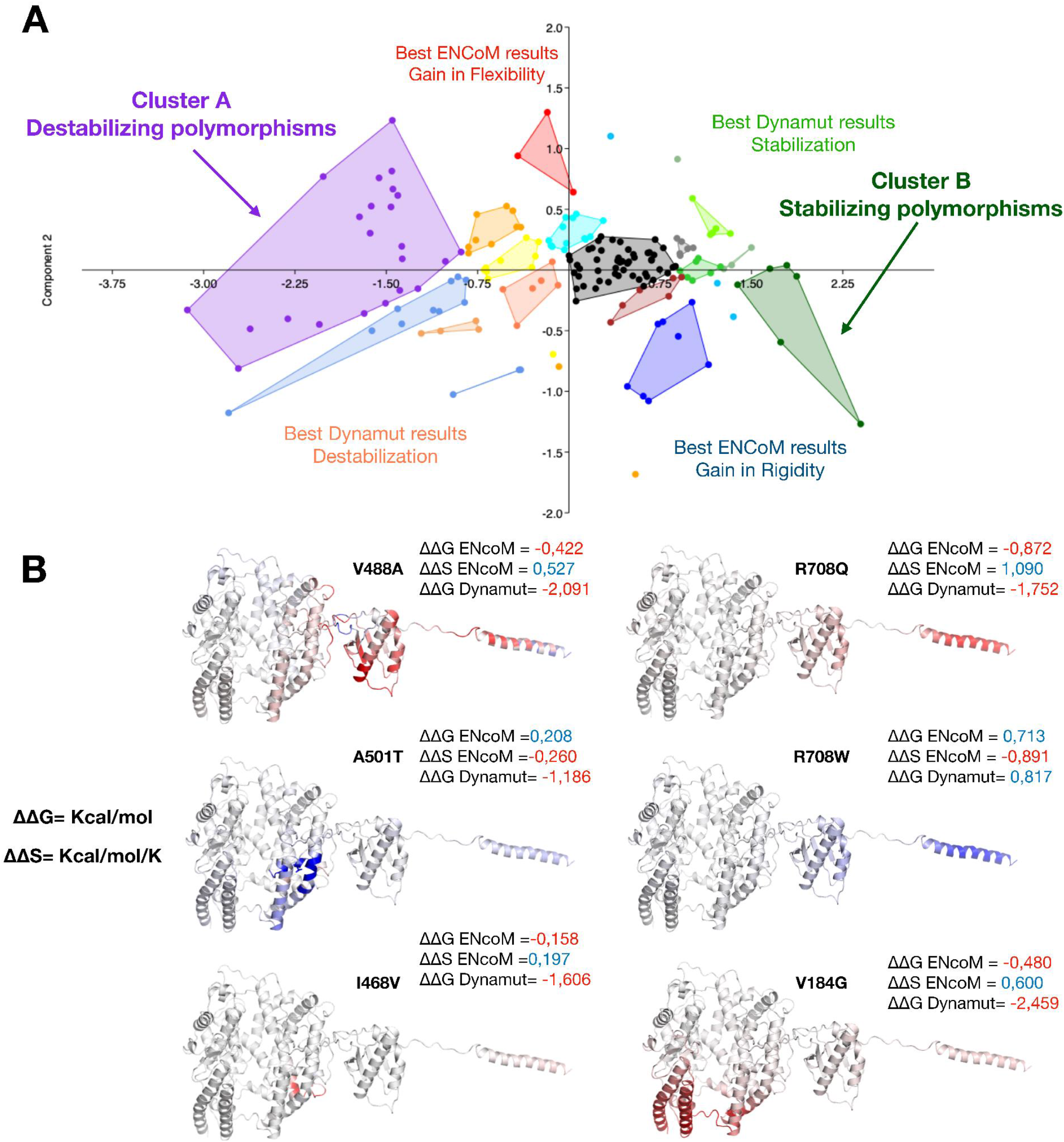
ACE2 SNPs analyzed by Dynamut and ENCoM. **a** Ordination plot (PCA) obtained with PAST to evaluate the results of Dynamut and ENCoM. This set of data includes all SNPs by dbSNP database, excluding SNPs in contact interfaces. **b** Effects on structure flexibility calculated as ΔΔG and ΔΔS variations between ACE2 and some polymorphic variants (red, gain in flexibility, blue, gain in rigidity). Illustrated SNPs were selected as the most common polymorphisms that gave the highest ΔΔG and ΔΔS variations.

All 197 amino acid residues that were reported as polymorphic in dbSNP were analyzed. In Supplementary Fig. 5a the ordination plots that were generated by clustering the effects of the single amino acid substitutions according to the Dynamut^36^ (left panels) and ENCoM^42^ (right panles) outputs are illustrated. The data of the amino acid substitutions in the three ACE2 interfaces (ACE2/SARS-CoV-2 Spike, ACE2/ACE2, ACE2/B0AT1) were then extrapolated (Supplementary Fig. 5b), and were excluded from the analysis that was aimed at predicting structural (indirect) effects of amino acid substitutions on binding interfaces. The resulting subset of data (Supplementary Fig. 5c) was combined by matching Dynamut^36^ and ENCoM^42^ predictions, and used to generate the final ordination plot shown in Fig. 6a. Results indicate that a considerable number of the amino acid substitutions are able to either stabilize or destabilize the binding interfaces with possible consequences on either ACE2 function and/or ACE2/SARS-CoV-2 Spike interaction. In particular, in Fig. 6b, the effects of amino acid substitutions on structure flexibility were calculated as ΔΔG and/or ΔΔS values comparing ACE2 and each polymorphic variant (red gain in flexibility, blue gain in rigidity). SNPs I468V, V488A and A501T were selected as the most common polymorphisms that gave the highest ΔΔG and ΔΔS values in our dynamic model.

## Discussion

Although with limitations and caveats of *in silico* technology, this study addresses the question of whether some ACE2 SNPs may be associated with a different individual susceptibility to COVID-19. To alleviate these limitations, we used a combination of bioinformatics tools, and tested different crystallographic models.

Four months after the spread of the SARS COVID-19, its worldwide distribution remains extremely uneven. Lethality is even more inhomogeneous among and within countries, with figures of 12.6% in Italy^43^, and 0.6% in South Korea^44^. Although differences in mortality might have various causes, including access and efficiency of health systems, total number of people tested, presence and severity of symptoms in tested populations, they are so impressive that it seems legitimate to search for other factors possibly related to individuals as the elements of a population. Ultimately, infectivity and lethality do not seem linearly related, and probably represent problems to be solved with different, albeit complementary, approaches.

Basic aspects of epidemiology of the disease warrant some considerations: differently from other countries, in South Korea (which adopted a policy of extensive PCR screening), women represent 63% of infected people^44^ as opposed to 50% in Italy^43^ (where the policy has been to test only severely symptomatic cases for a long time). Lethality figures in women were 0.4% and 8.7% in South Korea and Italy, respectively, as opposed to 1% and 16.4% in men. It could be speculated that women are probably more prone to infection but often present a less severe disease. Although higher incidence of cardiac, respiratory and metabolic co-morbidities are probably responsible for more severe form of infection in men, estrogen-induced upregulation of ACE2 expression would explain increased susceptibility of women to a less severe and often asymptomatic form of disease. Furthermore, the ACE2 gene is located on Xp22, in an area where genes are reported to escape from X-inactivation, further explaining higher expression in females^45,46^.

On the other hand, it has been hypothesized that, regardless of sex, pharmacological (antihypertensive drugs, such as ACE inhibitors and sartans) or environmental factors (NO2 pollution), capable of inducing an overexpression of ACE2 could be responsible of increased susceptibility to infection and/or greater severity^47^. ACE2 plays an essential role in the renin-angiotensin-aldosterone system, and its loss of function due to the massive binding of viral particles and internalization could constitute an essential element of the pathophysiology of pulmonary and cardiac damage during COVID-19 infection^47,48^. In this context it should be underlined that ACE2 probably plays a dual role in the dynamic of infection and disease course. While at beginning ACE2 overexpression may increase the entry of the virus into the cell and its replication, its consequent viral-induced loss of function results in an unopposed accumulation of angiotensin II that further aggravates the acute lung injury response to viral infection. Indeed, in the rodent blockade of the renin-angiotensin-aldosterone system limits the acute lung injury induced by the SARS-CoV-1 spike protein^49^, suggesting that if ACE2 function is preserved (because of increased baseline expression, as especially seen in pre-menopausal women), clinical course of infection might be less severe.

It has been suggested that polymorphisms in the ACE2 gene could reduce the spike affinity, with subsequent lower susceptibility to infection: in this hypothesis, their geographical / ethnical distribution could explain the strong discrepancies in infection rate and/or lethality observed worldwide^47^. Effectively, we showed by Network plot and Non-Metric Multidimensional Scaling that most of the SNPs diffused worldwide did not affect significantly the interaction of ACE2 with SARS-CoV-2 Spike. S19P was one of the rare polymorphisms able to potentially affect this interaction, by lowering the affinity. This polymorphism is more frequent in African populations, but its diffusion (0.3%) remains too low to explain, except in minimal part, the reduced death toll observed so far in that continent, and, more generally, the enormous differences in geographical spread of infection and lethality. However, it seems clear that the affinity of the virus for ACE2 is a key determinant of its infective potential: in order to choose the experimental model capable of reproducing the essential aspects of human infection, Chan and colleagues^50^ determined *in silico* the spike / ACE2 affinity in primates and in a series of experimental animals, observing that the binding energy is maximal in primates (−62.20 Rosetta energy units (REU)), intermediate in Syrian hamster (−49.96 REU), lower in bat (−39.60 REU). This allowed the authors to predict that hamsters could be infected, which was experimentally confirmed –underlining the reliability of *in silico* modeling- and could be subsequently at the origin of inter-animal transmission. However, hamsters, although developing clinical signs of the infection and relative histopathological changes, did not die^50^: we speculate that lethality may be related to Spike/ACE2 affinity. On the other hand, the lower affinity in bat could explain –besides a better immune control-why these animals are carriers without dying.

In the same study, Chan and colleagues^50^ showed that the binding energy between ACE2 and Spike of SARS-CoV, responsible for the 2002 epidemic, was −39.49 REU as compared to −58.18 of human ACE2. After that epidemic, attempts at developing mouse experimental models, resulted only in mild lung inflammation and rapid viral clearance, until development of transgenic mice expressing human ACE2 under regulation of a global promoter or cytokeratin 18 promoter, which developed rapidly lethal infection after intranasal viral inoculation^51,52^. Interestingly, intranasally CoV infected transgenic mice expressing human ACE2 driven by the mouse ACE2 promoter, developed severe disease (on clinical and histopathological grounds, including typical interstitial pneumonia and widespread extrapulmonary organ damage) without dying; furthermore their viral clearance was markedly slower as compared to wild type mice which had only mild abnormalities^53^.

So, if modestly SNP-determined lower affinity between spike and ACE2 does not seem to explain the differences in the distribution and lethality of the disease in humans, we hypothesize that the question can be addressed in a specular way: perhaps polymorphisms responsible for higher affinity can be responsible of higher severity of disease, especially when very high affinity receptors are overexpressed because of the above mentioned environmental and pharmacological factors. Obviously underlying diseases would contribute to an even more severe course of the disease, with an intense viral replication capable of infecting in turn a large number of persons, including some individuals with similar ACE2 polymorphisms, and so on. Our *in silico* models allowed us to identify K26R and K26E as SNPs with a possible increase in Spike/ACE2 affinity. K26R SNP is relatively frequent in European people with a frequency about 0.5%, which would correspond to a potential target population of 2,230,000 people at the European Union level.

In addition to FireDock^34, 35^ simulations that led to predict the possible effects of S19P, K26R and K26E ACE2 SNPs, Dynamut^36^ and ENCoM^42^ tools were used to compare dynamic features of ACE2 and its polymorphic variants in order to analyze the possible indirect effects on binding interfaces of SNPs that are located outside these interfaces. SNPs I468V, V488A and A501T were identified as the most common SNPs that may produce these indirect effects in dynamic models. Although the precise effects of these SNPs on the interaction between ACE2 and SARS-CoV-2 or SARS-CoV Spike proteins have to be determined in more detail, nevertheless, it is desirable to use dynamic modeling to unmask indirect effects of SNPs.

It seems necessary to confirm *in vivo* that, among patients with serious disease and/or fatal outcome, polymorphisms responsible for a very high Spike/ACE2 affinity are more frequent than among patients with less severe/asymptomatic disease or even than in general population. Obviously, the impact of these polymorphisms on severity of outcome should be weighted by appropriate demographic and clinical factors. If these differences were confirmed, this would pave the way for the identification, on a population scale, of healthy individuals whose molecular phenotypes would be responsible for more serious disease. Apart from the usual social distancing measures, which could be reinforced for these cases, targeted drug prevention strategies could be evaluated. It could be logical to assess pharmacological prophylactic interventions, as proposed in categories of healthy people at particular risk of exposure such as care-givers. In particular, chloroquine, interfering with N-terminal glycosylation of ACE2, could lower its affinity for spike, thus representing an interesting candidate. In our *in silico* model, we found that removal of glycosidic moieties weakened the interaction between SARS-CoV-2 spike and ACE2. The serine protease inhibitor camostat mesylate, approved in Japan to treat unrelated diseases, has been shown to block TMPRSS2 activity^54,55^ and is thus another interesting candidate.

On the other hand, the identification of broader categories of people with lower risk of developing severe disease, could allow a safer exit from the lock-down phase, while facilitating the establishment of a faster herd immunity, and waiting reliable serological tests and, above all, effective vaccines.

On the basis of our *in silico* study we speculate that infection and mortality are determined at individual level by different factors including the amount of expression of ACE2 and its affinity with the spike protein. While the level of ACE2 expression possibly determinates the probability of infection in the presence of an adequate inoculum, the severity of the disease is mainly determined by the phenotype affinity for the spike protein. Clinical studies are urgently required to confirm the present mechanistic hypothesis.

## Materials and Methods

### Databases

3D structures of proteins were downloaded from PDB (RCSB Protein Data Bank^56^). We focused our analysis on 2AJF for SARS-CoV^27^ (DOI: 10.2210/pdb2AJF/pdb)and 6VW1^30^ (DOI:10.2210/pdb6VW1/pdb), 6M17^20, 21^ (DOI:10.2210/pdb6M17/pdb) 6LZG^28^ (10.2210/pdb6LZG/pdb), 6M0J^29^ (10.2210/pdb6M0J/pdb)models for SARS-COV-2. dpSNP database ^57,58^ was used to identity the ACE2 receptor SNPs, and to select the most diffused ones. Functional information was acquired by UNIPROT database^59^. Chimera^60^ was used as a tool for Image generation, 3D mapping, PDB managing and to analyze the results.

### Binding interface characterization

The selected PDB models were analyzed by a structural point of view using Chimera software in order to identify the glycosylation sites and the secondary structures of proteins involved in the binding between ACE2 and Spike protein receptor binding domain (RBD). To estimate the effect of glycosylations we implemented a static model. Chimera was used to remove glycosydic residues, while FireDock^34,35^ was used to compute the global energy scores between the native structures and the de-glycosylated models. On the other side, starting from the entire list of SNPs, SSIPe (EVOEF)^32^ was used to identify the residues involved in the binding interfaces. A second step, which was carried out with SSIPe (SSIPe)^33^, was aimed at estimating the effects of single SNPs, and to generate mutant models. Different SNPs lists were obtained, which were compared, and used to identify the most stable binding amino acid residues. SSIPe analysis performed with the PDB model 6M17 was used to map: the ACE2/Spike protein interaction interface, ACE2 /ACE2 dimerization interface, and B0AT1/ACE2 interaction interface. The model contains ACE2 in the dimeric form (with the hydrophobic domains) and B0AT1, while in all the others models the transmembrane domains, ACE2 /ACE2 interface and B0AT1/ACE2 interface are absent.

### Dynamic analysis of ACE2 structure

To obtain a dynamic model of the ACE2 we used different tools. Dynamut^36^ was used to calculate the general dynamic features of the ACE2 in 6M17 model leading to the identification of two domains characterized by high deformations and high fluctuations scores, respectively. To validate these results and to compare them with Spike proteins fluctuations, we used CABSflex^41^ to analyze Spike protein chains in 2AJF and 6VW1 models, and ACE2 chain in 6M17 model. Another Dynamut analysis was performed on 6VW1 model, considering only the structure of the globular head of ACE2. The transmembrane domain was mapped on a 3D file using TMHMM to predict hydrophobic helix. A molecular dynamics approach was used to model the behavior of the transmembrane domain. CHARMM-GUI^37,38^ was used to implement the model of a DPPC (phosphatide) bilayer that embedded ACE2, while NAMD^40^ was used to perform the analysis via VMD (NAMD GUI)^39,40^. Chimera was used to select the frames, and to measure the oscillations of interface residues.

### Effect of mutations on ACE2 dynamic features

Dynamut^36^ was used to estimate the effect of SNPs on structural dynamics features. This tool generated two results with the two algorithms ENCoM^41^ and Dynamut. A quartile-based clustering was performed with 197 SNPs obtaining two equivalent plots where results were clustered in two ways: ENCoM groups and Dynamut groups. After this step, we extrapolated data related to the three interfaces (ACE2/Spike protein, ACE2/B0AT1 and ACE2/ACE2), because the analyzed SNPs can directly interfere on either binding to Spike proteins, or ACE2 homo-dimerization or ACE2/B0AT1 binding. In order to select the best results from this last subset, the clusters for Dynamut and for ENCoM were combined. PCA plots were generated using PAST software^61^.

### Docking: dynamic models of ACE2 / Spike protein interactions

To obtain the protein complex models with and without glycosylations we used two software: PathDock^62^ and Gramm-X^63^, respectively. Resulting complexes were submitted to FireDock^34,35^ to evaluate the energy score. As receptors and ligands we used all chains of ACE2 in PDB models (2AJF, 6VW1, 6LZG, 6M0J, 6M17), and the models of isoform X1 and X3, which were generated by using SwissModel^64^ tool, while all chains of the Spike protein in the PDB files were used as ligands. For each software, the total simulations number was 35. All the obtained results were screened excluding: i) the high-energy complexes, ii) bad-orientated solutions (overlapping of receptor and ligand chain) and iii) off-target solutions (binding in membrane helix, in B0AT1 interface or ACE2 / ACE2 homodimerization interface). The selected results were used as inputs to superposition using: the docking solutions as a model, the chains of B0AT1(2X) / ACE2 (dimeric) from 6M17 model, and Spike protein trimeric structure, which were obtained by using I-Tasser^65^ (https://zhanglab.ccmb.med.umich.edu/COVID-19/). To obtain multiple conformations we performed more than one superposition (Chimera) on the solutions changing the aligned chains. The final results were analyzed using Chimera.

### African and European static and dynamic models

To gain information about the geographical distribution and abundance of ACE2 SNPs, we analyzed dbSNP^57,58^ and available databases: GnomAD-Exomes, TopMed, ExAC, GnomAD-Genomes, GO Exome Sequencing Project, 1000Genomes. The frequency values of the most abundant SNPs (1KGB database) were reported in Supplementary Table S4. Data were analyzed by PAST^61^ that generated NM-MDS ordination plots. Additionally, network and between-database correlation analyses (PAST^61^) were carried out in order to clarify the relationship between SNPs, and the correlation between data of all databases.

To model the binding of S19P ACE2 we implemented two types of analyses: static analysis and dynamic analysis. The models with substitutions of one single amino acid were selected form SIPPe results (static model), and were submitted to FireDock^34, 35^ server. In this step we focused on mutant and wild type models of 6VW1 and 2AJF. Serine in position 19 is the first residue of the mature chain of ACE2, while residues from 1 to 18 form a signal peptide. To prove if this SNP changes the cut site of the ACE2 precursor we used Signal IP 5.0^66^. The sequence of this predicted mutant was submitted to I-Tasser server^65^ in order to obtain the mutant models. Using this predicted model as a receptor and SIPPe complexes as a reference we superposed the structures obtaining two static models (one for ligand: 2AJF and 6VW1). The dynamic models were implemented using Gramm-X (ligands Spike chains of 6VW1 and 2AJF; receptor mutant models). All models obtained were submitted to FireDock^34, 35^ in order to obtain a value of global energy.

To model the binding K26R or K26E ACE2 we used a set of static models. Firstly, starting from the dataset of the mutant models selected by SIPPe^33^ results, we calculated changes in global energy. This step was repeated for three models: 6VW1, 2AJF and 6M17. For this last model, we calculated the variation in terms of global energy in two conformations, using as reference models both: i) ACE2-spike chains B-E and ii) ACE2-Spike chains D-F. In order to estimate the effect of different ligands, we used the models 6VW1 form SIPPe as reference structure and receptor, and all Spike models reported in this study as ligands. In a similar manner, other static models from SSIPe were used as models to estimate the variation in terms of free energy related to polymorphisms that map on the ACE2 dimerization interface and ACE2-B0AT binding interface.

## Acknowledgements

We wish to thank prof. Diane Damotte (University of Paris) for advice and critical reading of the manuscript.

## Author contributions

M.A., P.A.: conception, coordination, designing, writing.

M.C.: experimental set-up, pipeline development, *in silico* analysis;

P.F., A.I., designing, data providing;

## Competing interests

The authors declare no competing interests.

## Materials & Correspondence

The authors declare no competing interests.

## Legends to Supplementary Figures

**Supplementary Fig. 1.**
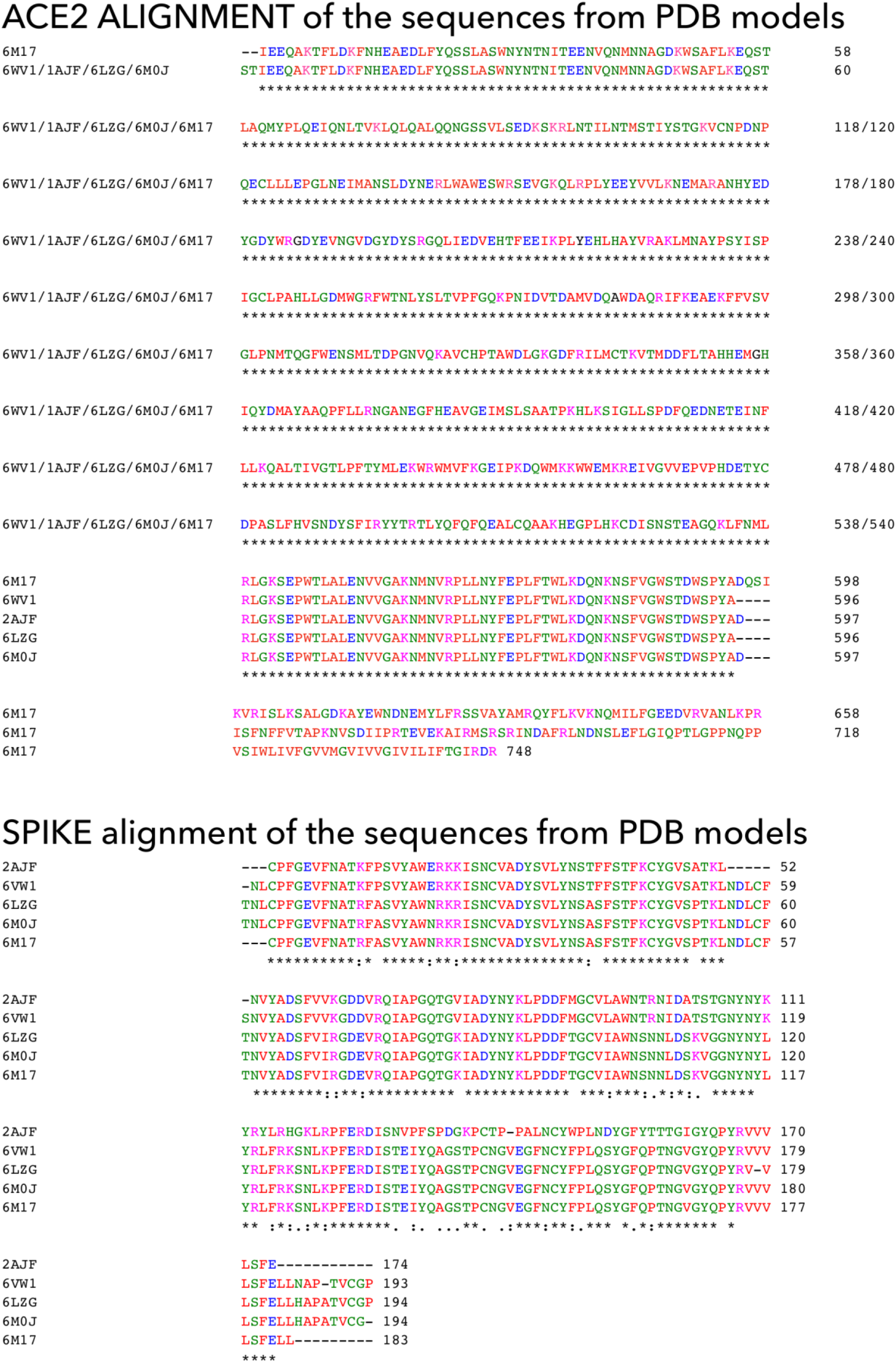
ClustalO alignments. ClustalO alignments of human ACE2 amino acid sequences, and SARS-CoV, SARS-CoV-2 and chimeric SARS-CoV/SARS-CoV-2 Spike RBDs used for X-ray cristallography models

**Supplementary Fig. 2.**
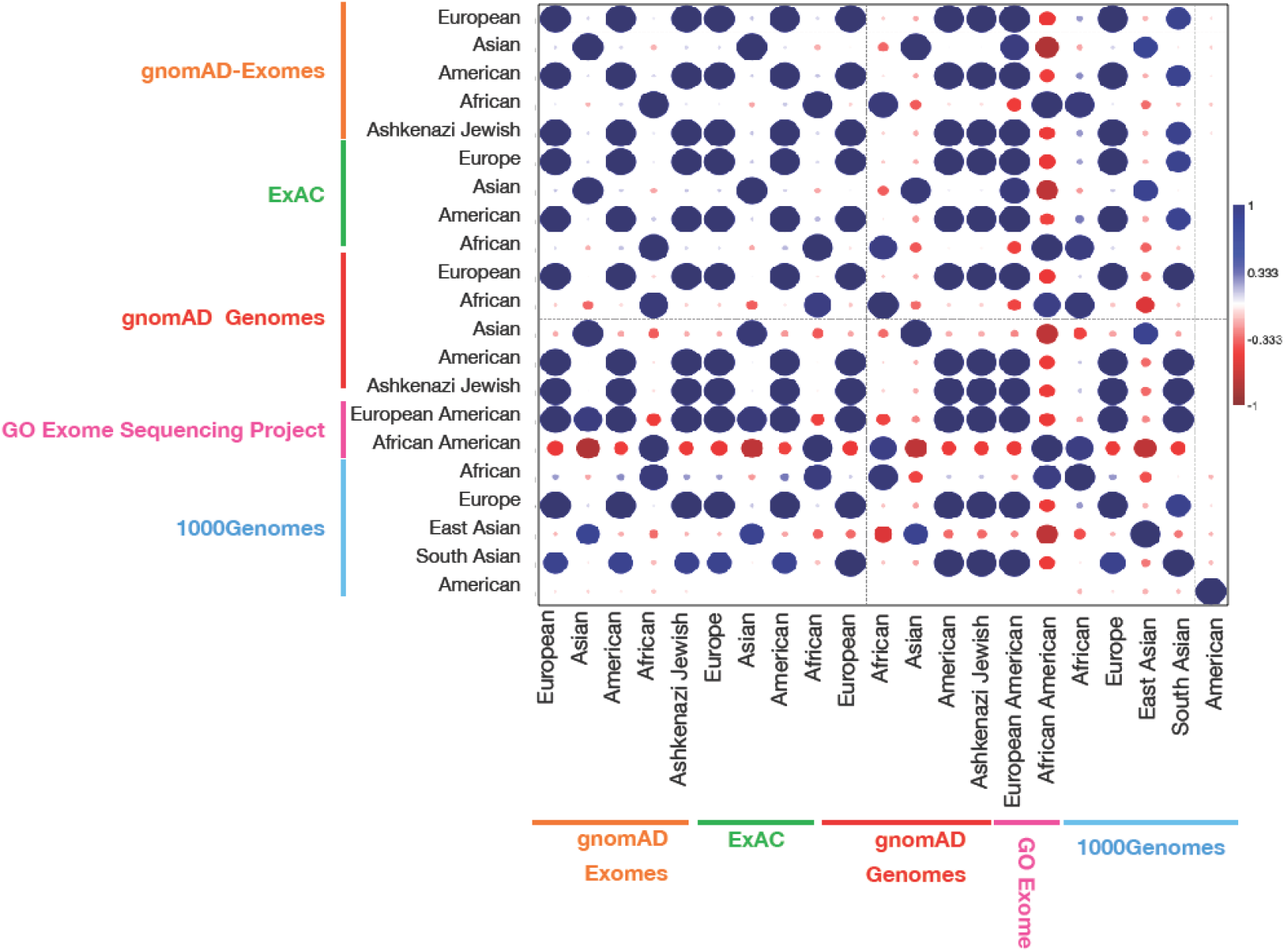
ACE2 SNPs data from different databases. Correlation analysis of ACE2 SNPs data from different databases is illustrated.

**Supplementary Fig. 3.**
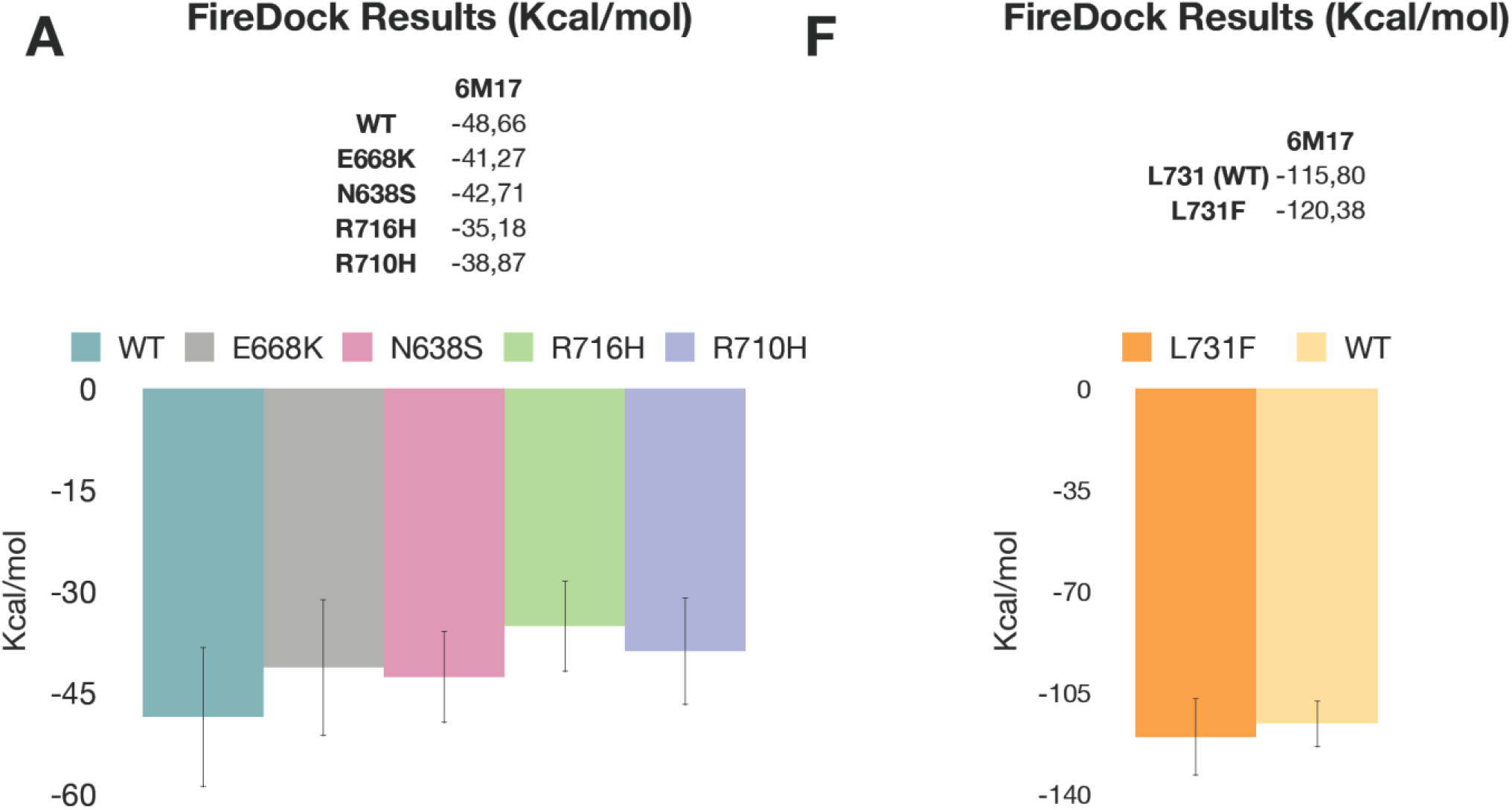
Selected ACE2 SNPs affecting binding interfaces. FireDock results predicting the effects of several ACE2 amino acid replacements on ACE2/ACE2 (left) and ACE2/B0AT1 (right) interaction.

**Supplementary Fig. 4.**
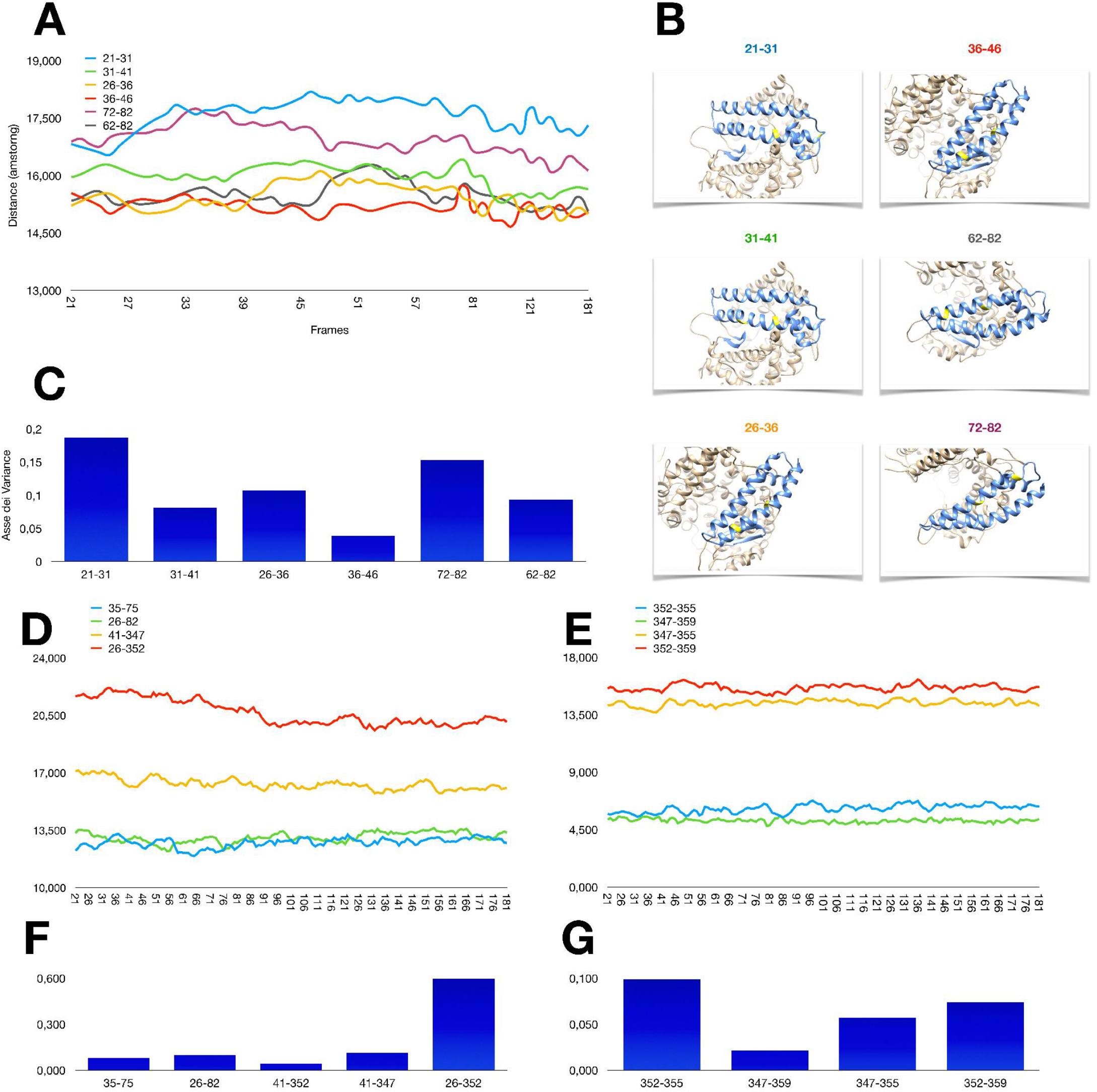
Oscillation of amino acid residues in ACE2 binding interfaces. **a** Oscillation plot of the distance between ACE2 amino acid residues mapping in the two helices that are involved in binding with Spike proteins. **b** 3D images of regions containing the amino acid residues shown in panel **a**. **c** Variance of the distance between ACE2 amino acid residues mapping in the two helices that are involved in binding with Spike proteins. **d-e** Oscillation plot of the distance between the two helices and beta-sheet (**d**), and between the residues of the beta-sheet (**e**). **f-g** Variance of the distance of amino acid residues between the two helices and beta-sheet (**f**), and between the residues of the beta-sheet (**g**).

**Supplementary Fig. 5.**
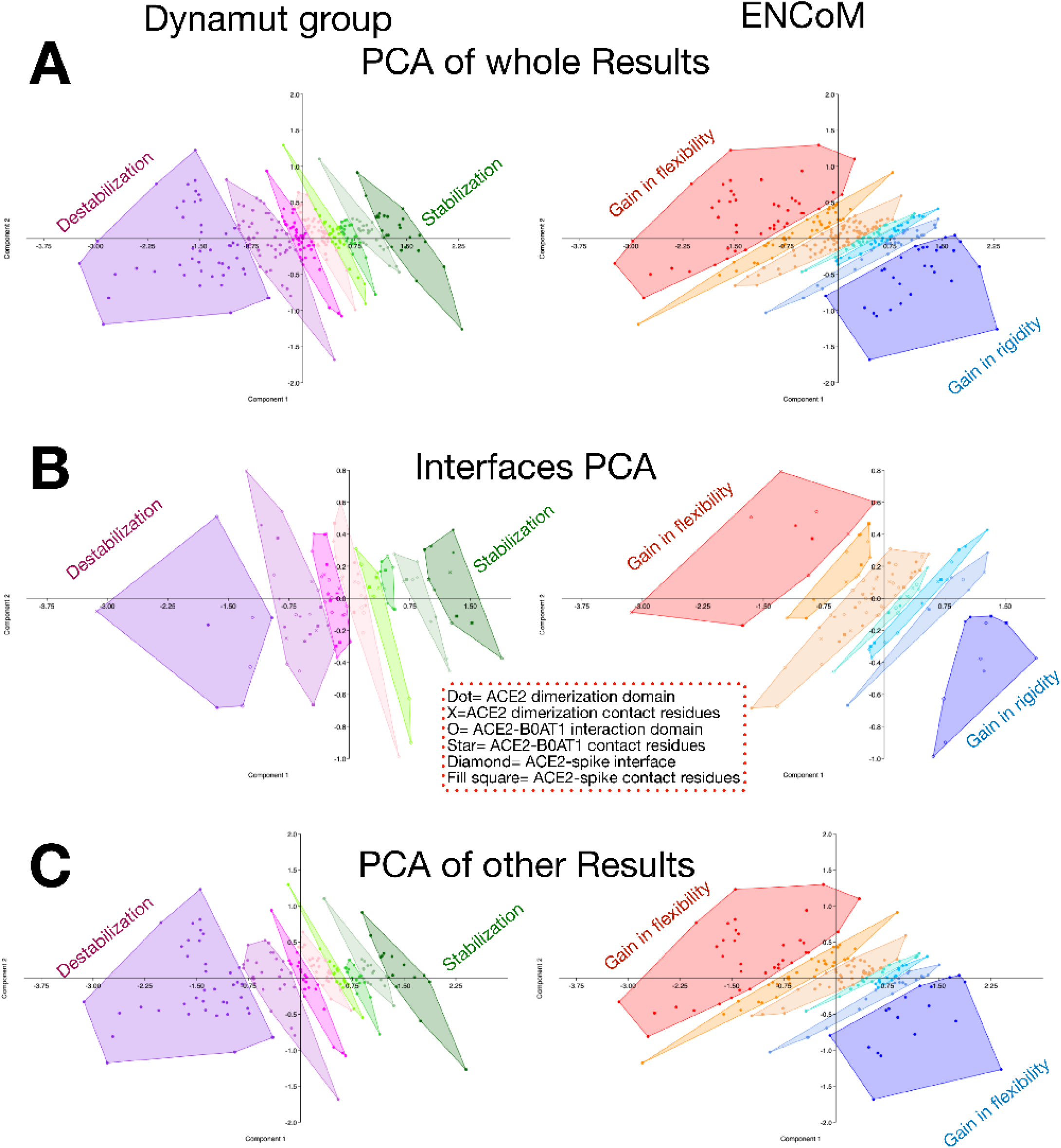
Ordination plots of Dynamut and ENCoM results. **a** Ordination plot (PCA) obtained using PAST to evaluate the results of Dynamut (left panels) and ENCoM (right panles) outputs. Clustering involved 197 SNPs and was based on ΔΔG and/or ΔΔS values comparing ACE2 and each polymorphic variant. **b** Ordination plot (PCA) showing the results of SNPs mapping within the ACE2 interfaces (ACE2/SARS-CoV-2 Spike, ACE2/ACE2, ACE2/B0AT1). C) Ordination plot (PCA) showing the results of SNPs Ordination plot (PCA) showing the results of SNPs that mapping outside the ACE2 interfaces.

## Supplementary Tables (Excel File)

**Supplementary Table 1. ACE2 PDB models and glycosylation.**

**Supplementary Table 2. EVOEF results.**

**Supplementary Table 3. SSIPe results.**

**Supplementary Table 4. ACE2 SNPs metadata.**

**Supplementary Table 5. GrammX-FireDock results.**

**Supplementary Table 6. PatchDock-FireDock results.**

**Supplementary Table 7. ENCoM and Dynamut results.**

## Notes

### Competing Interest Statement

The authors have declared no competing interest.

